# Colistin resistance prevalence in *Escherichia coli* from domestic animals in intensive breeding farms of Jiangsu Province, China

**DOI:** 10.1101/307983

**Authors:** X. Zhang, B. Zhang, Z. Yu, Y. Guo, J. Wang, P. Zhao, J. Liu, K. He

## Abstract

The global dissemination of colistin resistance has received a great deal of attention. Recently, the plasmid-mediated colistin resistance encoded by *mcr*-1 and *mcr*-2 genes in *Escherichia coli* (*E.coli*) strains from animals, food, and patients in China have been reported continuously. To make clear the colisin resistance and *mcr* gene spread in domestic animals in Jiangsu Province, we collected fecael swabs from pigs, chicken and cattle at different age distributed in intensive feeding farms. The selected chromogenic agar and *mcr*-PCR were used to screen the colisin resistance and *mcr* gene carriage. Colistin resistant *E.coli* colonies were identified from 54.25 % (440/811) pig faecal swabs, from 35.96 % (443/1232) chicken faecal swabs, and 26.92 % (42/156) from cattle faecal swabs. Of all the colisin resistant *E.coli* colonies, the positive amplifications of *mcr*-1 were significantly higher than *mcr*-2. The *mcr*-1 prevalence was 68.86 % (303/440) in pigs, 87.58 % (388/443) in chicken, and 71.43 % (30/42), compared with 46.82 % (206/440) in pigs, 14.90 % (66/443) in chicken, and 19.05 % (8/42) in cattle of prevalence of *mcr*-2. Co-occurrence of *mcr*-1 and *mcr*-2 was identified in 20 % (88/440) in pigs, 7.22 % (32/443) in chickens, and in 9.52 % (4/42) cattle. These data indicate that *mcr* was the most important colistin resistance mechanism. Interventions and alternative options are necessary to minimise further dissemination of *mcr* between food-producing animals and human.

**IMPORTANCE:** Colistin is recognized one of the last defence lines for the treatment of highly resistant bacteria, but the emergence of resistance that conferred by a transferable plasmid-mediated *mcr* genes to this vital antibiotic is extremely disturbing. Here, we used *E. coli* as an index to monitor drug resistance in domestic animals (pigs, chicken and cattle). It was found that the colistin resistance widely occurred at all ages of domestic animals and the *mcr*-dependent mechanism dominated in *E.coli*. We also found that the elder and adult animals were a reservoir of resistant strains, suggesting a potential food safety issue and greater public health problems.

## 1 INTRODUCTUON

Colistin is recognized one of the last defence lines for the treatment of highly resistant bacteria, but the emergence of resistance that conferred by a transferable plasmid-mediated *mcr*-1 gene to this vital antibiotic is extremely disturbing. Actually, the mechanism of colistin resistance can be generally classified as *mcr*-independent or *mcr*-dependent. In a Morbidity and Mortality Weekly Report (MMWR) in September 2016, Vasquez and colleagues isolated a shiga-toxin-producing *Escherichia coli* (STEC) O157 with the *mcr*-1 gene in the whole genome sequence from stool [1]. In November 2016 in *the Lancet Infectious Diseases*, Liu et al. reported finding a transferable plasmid-mediated *mcr*-1 gene in *E. coli* isolates from animal food in China [2]. Compared with *Klebsiella pneumoniae* and *Pseudomonas aeruginosa*, in *E. coli* rare colistin resistance was mediated by chromosomal mutations and possibly imposed a fitness cost to the organism [3], which suggested that *mcr*-dependent colistin resistance perhaps was the major mechanism in *E.coli*, and would promote colistin resistance transmission among bacteria by plasmid transfer and chromosomal recombination. In China, [4]since the early 1980s colistin has been used in animals as a therapeutic drug and feed additive, which emphasizes that the use of colistin in animal feed has probably accelerated the dissemination of *mcr* gene in animals and subsequently humans.

## 2 MATERIALS AND METHODS

### 2.1 Sample collection

From March 2015 to December 2016, a surveillance of colistin resistant *E.coli* was conducted in Jiangsu Province, China. A total of 2199 faecal swab samples (**Table 1**) were collected from pigs, chicken and cattle. 811 faecal swab samples were collected from suckling piglets, weaned piglets, fattening pigs, and sows. 1232 faecal swab samples were collected from chicks, egg-laying growers and laying hens. 156 faecal samples were collected from calves, growing cows and milking cows.

**Table 1:**
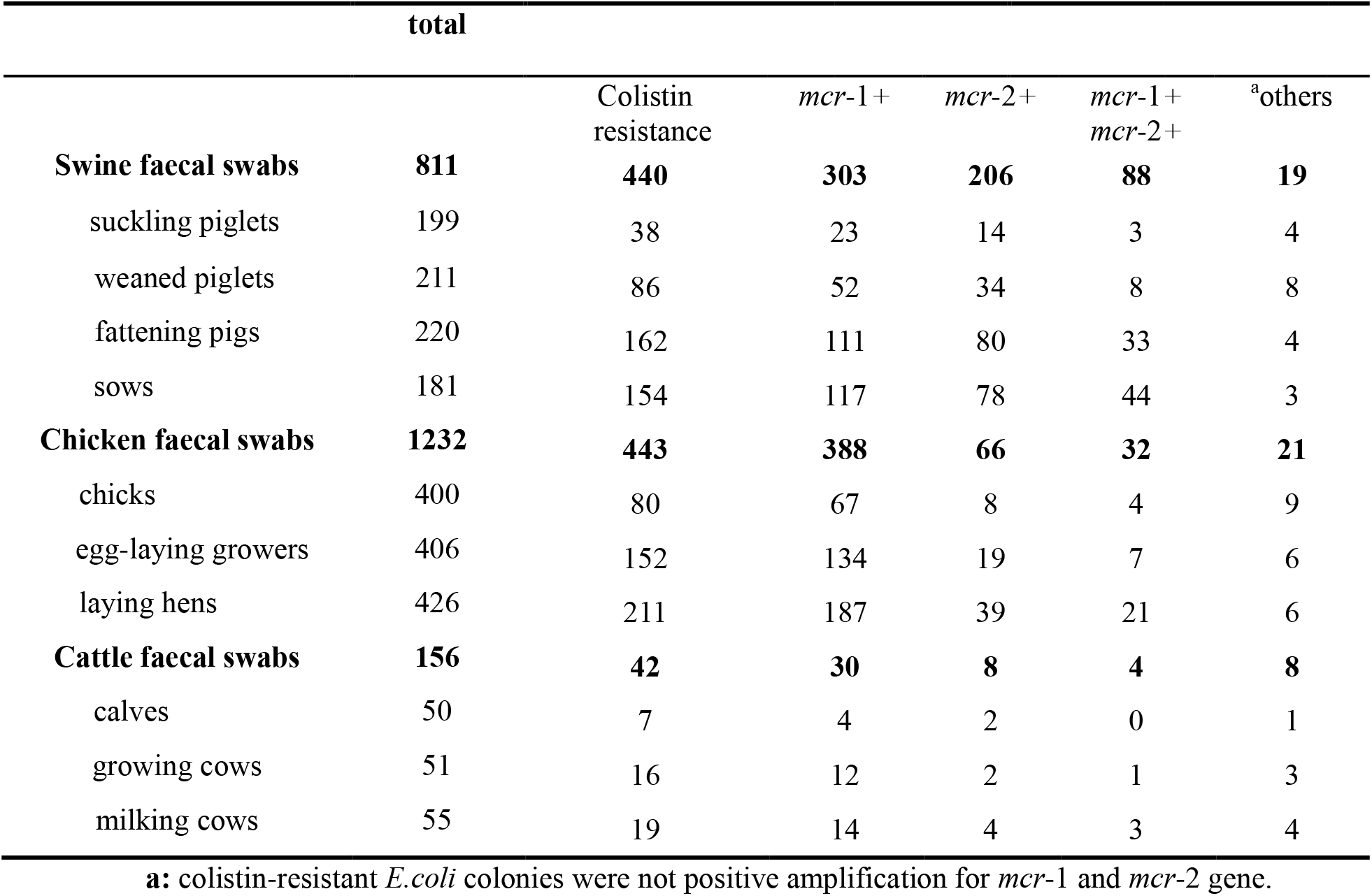
Categories of samples and numbers of colistin resistance-positive *E. coli*. and *mcr*-positive *E. coli*. in this study.

### 2.2 Colistin resistance screening

*E. coli* has been identified as an index for monitoring drug resistance [5–6]. Here, we used *E.coli* selected chromogenic agar with 10 μg/mL of colistin sulphate [1] to test drug resistance to *E.coli* in domestic animal faeces. Each swab was dipped in 2 mL PBS for two hours at 4°C, and then homogenised by vortex. The homogenates were centrifuged at 500 rpm for 15 minutes. After the aspirated supernatants were centrifuged at 12,000 rpm for 5 min, the pellets were suspended with 1 mL PBS. Tenfold dilution series of 100 μL of the suspended pellets were plated onto *E.coli* selected chromogenic agar (HopeBio Biotech Corp., China) containing colistin sulphate. After overnight incubation at 37°C, the blue-green ones were counted as *E.coli* colonies (HopeBio Biotech Corp., China). If necessary, the faecal swabs were dropped into Tryptic Soy Broth (TSB) with antibiotics for enrichment and then bacterial culture was plated onto *E.coli* selected chromogenic agar (HopeBio Biotech Corp., China).

### 2.3 *mcr*-1 and *mcr*-2 screening

All blue-green colonies were picked into Luria-Bertani (LB) broth for 6 h enrichment, and bacterial culture were prepared DNA template by conventional boiling method. For the *E.coli* colonies identified, PCR was used to verify them by primer pairs of P1-F and P1-R [7] from the 16sRNA gene. For *mcr*-1 gene and *mcr*-2 screening, two primers pairs of P2-F/R [1] and P3-F/R [8] were used to amplify them. All the positive amplifications were sequenced by Genscript Corporation (Nanjing, China). Primers used in this study were listed in **Table 2**.

**Table 2:**
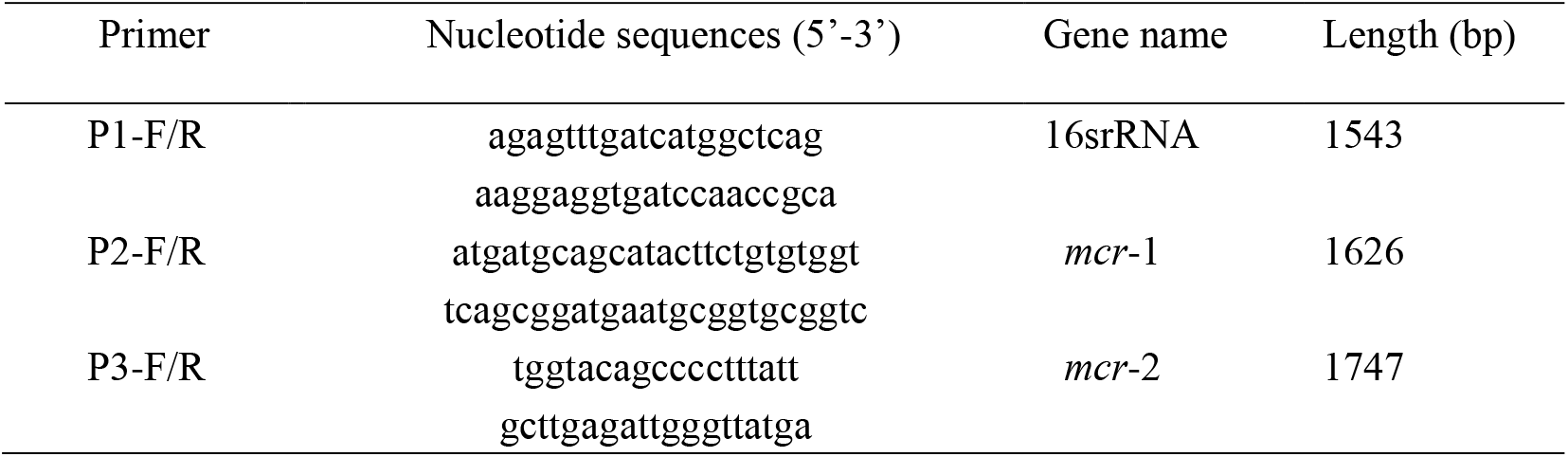
Primers used in this study

## 3. RESULT

### 3.1 Plate screening for colistin-resistant *E.coli* colonies

All blue-green colonies from *E.coli* selected chromogenic agar were recognized *E.coli* colonies after double identification using primer pairs of 16sRNA. In pigs, colistin resistant colonies were identified from 19.10 % (38/199) of sucking piglet, 40.76 % (86/211) of weaned piglet, 73.64 % (162/220) of fattening pig, and 85.08 % (154/181) of sow. In chicken, colistin resistant colonies were identified from 20 % (80/400) of chick, 37.44 % (152/406) of egg-laying grower, and 49.53 % (211/426) of laying hen. In cattle, colistin resistant colonies were identified from 14 % (7/50) of calve, 31.37 % (16/51) of growing cow, and 34.55% (19/55) of milking cow. Data on prevalence of colistin resistance in swab samples from all ages of domestic animals are presented in **Fig 1**.

**Fig. 1.**
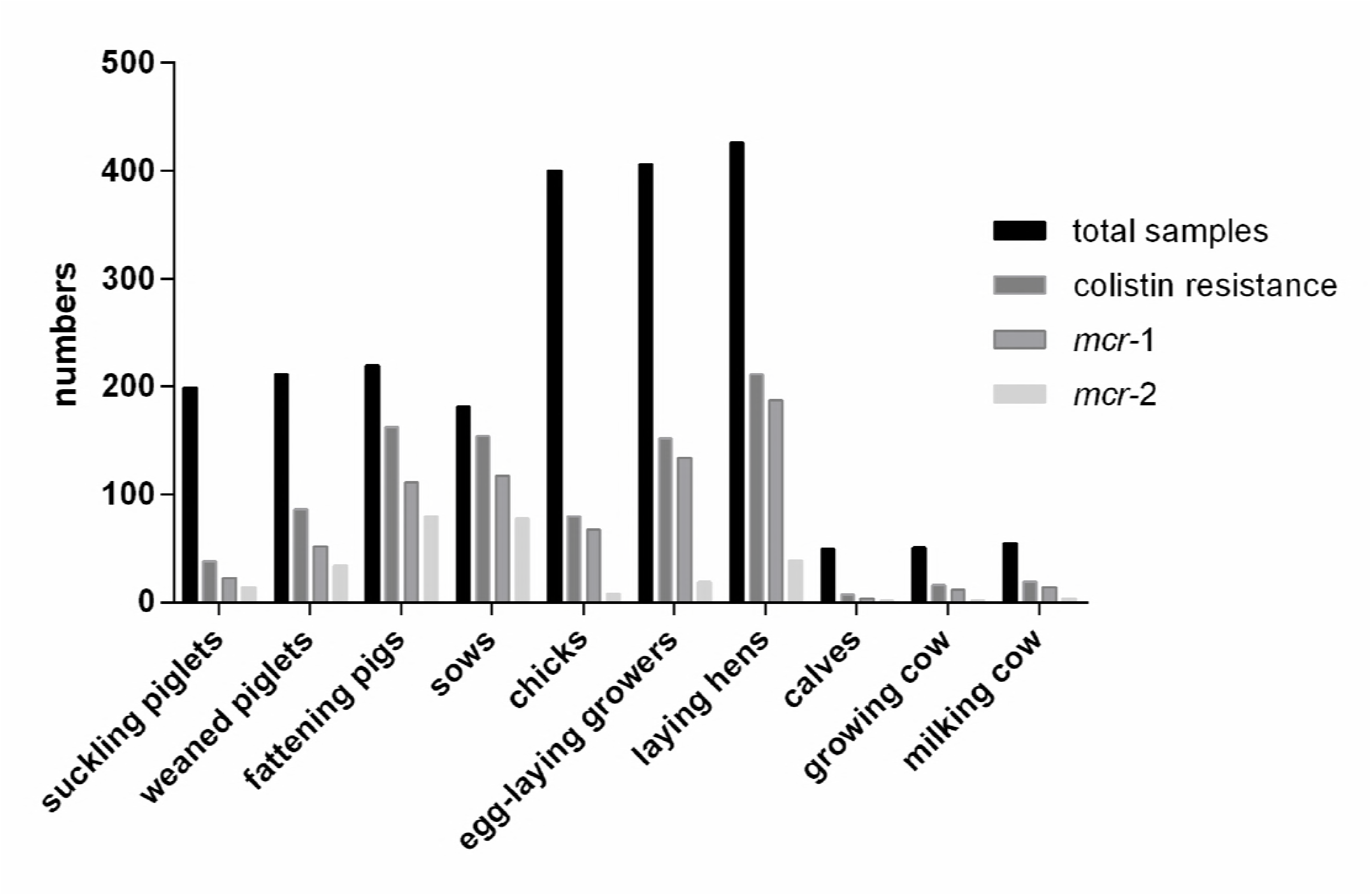
Prevalence of colistin resistance and *mcr* genes in all ages of animals

### 3.2 Prevalence of *mcr*-1

The *mcr*-1 was identified in colistin-resistant *E.coli* colonies from all ages of pigs, chickens, and cattle. The *mcr*-1 prevalence was 68.86 % (303/440) in pigs, 87.58 % (388/443) in chicken, and 71.43 % (30/42) in cattle (**Fig. 1**). For pigs, the specific *mcr*-1 PCR identified the gene in 60.53 % (23/38) of suckling piglets, 60.47 % (52/86) of weaned piglets, 68.52 % (111/162) of fattening pigs, and 75.97 % (117/154) of sows. For chickens, the specific *mcr*-1 PCR identified the gene in 83.75 % (67/80) of chicks, 88.16 % (134/152) of egg-laying growers, and 88.63 % (187/211) of laying hens. For cattle, the specific *mcr*-1 PCR identified the gene in 57.14 % (4/7) of calves, 75.00 % (12/16) of growing cows, and 73.68 % (14/19) of milking cows.

### 3.2 Prevalence of *mcr*-2

The *mcr*-2 was identified in colistin-resistant *E.coli* colonies from all ages of pigs, chickens, and cattle. The *mcr*-2 prevalence was 46.82 % (206/440) in pigs, 14.90 % (66/443) in chicken, and 19.05 % (8/42) in cattle (**Fig. 1**). For pigs, the specific *mcr*-2 gene was amplified from 36.84 % (14/38) of suckling piglets, 39.53 % (34/86) of weaned piglets, 49.38 % (80/162) of fattening pigs, and 50.65 % (78/154) of sows. For chickens, the specific *mcr*-2 gene was amplified from 10 % (8/80) of chicks, 12.50 % (19/152) of egg-laying growers, and 18.48 % (39/211) of laying hens. For cattle, the specific *mcr*-2 gene was amplified from 28.57 % (2/7) of calves, 12.50 % (2/16) of growing cows, and 21.05 % (4/19) of milking cows.

### 3.3 Co-occurrence of *mcr*-1 and *mcr*-2

Both *mcr*-1 and *mcr*-2 positive amplifications were 20 % (88/440) in pigs, 7.22 % (32/443) in chickens, and 9.52 % (4/42) in cattle (**Table 1**). Dual positivity was identified in 7.89 % (3/38) of suckling piglets, 9.30 % (8/86) of weaned piglets, 20.37 % (33/162) of fattening pigs, 28.57 % (44/154) of sows, 5.00 % (4/80) of chicks, 4.61% (7/152) of egg-laying growers, 9.95 % (21/211) of laying hens, 6.25 % (1/16) of growing cows, and 15.79 % (3/19) of milking cows, but not in calves.

## 4. DISCUSSION

In the 1960s colistin was introduced into in food animal production in several countries for growth promotion, therapeutical and prophylactical purposes to control of *Enterobacteriaceae* infections, particularly for those caused by *E.coli* [5–6]. In 2016, Chinese scholars first reported that plasmid-mediated colistin resistance was encoded by the *mcr*-1 gene [1]. With this discovery, the higher prevalence of samples harboring *mcr*-1 gene in animal isolates compared to other origins raised alarm bell about the impact of colistin use on colistin resistance spread in animal production, livestock and poultry have been recognized as the major reservoir for colistin resistance transmission and amplification [9].

During 2015-2016, we collected 2199 faecal swabs from pigs, chicken and cattle to make clear prevalence of colisitn resistance in intensive breeding farms of Jiangsu Province. Our study using selected chromogenic agar with colistin showed that *E.coli* resistance to colistin occurred widely in pigs (54.25 %), poultry (35.96 %) and cattle (26.92 %), suggesting that colistin resistance was considerably serious, especially in pigs. From 2013 to 2014, it was reported that a high frequency of colistin resistance in *E. coli* from pigs (26.5%), from chickens (14.0%), and from cattle (0.9%) on farms in different geographic areas of China, including Jiangsu Province [10]. Increasing use of colistin in fodder in recent years may be the reason of the high prevalence of colistin resistance in these food animals. Here, in 811 pig samples, colistin resistant colonies were identified from 85.08 % (154/181) of sows and 73.64 % (162/220) of fattening pigs, significantly higher than 19.10 % (38/199) of sucking piglet, 40.76 % (86/211) of weaned piglets. The same patterns also were found in chicken 1232 samples and 156 cattle samples. The highest proportions of resistant *E.coli* colonies were identified from the adult animals, implying that the long-term selective pressure resulted in not only the highest prevalence of colistin resistance among *E. coli* isolates from adult animals found in this study, but also bacterial evolution and adaption from the piglet groups to adult groups [11]. Compared with the isolates from pigs and chickens recovered during 2013-2014, *E. coli* isolates collected during 2007-2008 (5.5%) and 2010-2011 (12.4%) showed significantly lower frequency of colistin resistance [12]. A high frequency of colistin resistance in *E. coli* from pigs on farm (24.1%) and from chickens on farm (14.0%) led to a high prevalence of colistin at pig slaughter (24.3%) and chicken slaughter (9.5%) in 2013-2014 [12]. The adult animals generally entered the slaughter house and the food chains, drug-resistant strains inevitably invaded our dining table for consumers to cause public health events. Sows are the reservoir of resistant strains, they give not only life to piglets, but also resistant strains to them, which promote drug resistance circulation among Chinese farms [13]. The link between animals and humans in terms of colistin resistant *E. coli* strain transfer following direct contact has recently been confirmed [14]. The overuse of antibiotics will promote the unrestricted expansion and circulation of drug-resistant strains among human-animals-environment.

While colistin is a last-line antibiotic used to treat multidrug resistant Gram-negative bacteria (GNB) isolated from food animals, raw meat, and humans in several countries [15], its efficacy is being compromised by the detected mobile colistin resistance genes, *mcr-1* at the end of 2015 [2], and subsequently *mcr*-2, *mcr*-3, *mcr*-4, *mcr*-5[8, 16]. Of all the colisin resistant *E.coli* colonies in our study, the *mcr*-1 was the predominant gene for the colistin resistance of *E.coli*, higher than *mcr*-2. The *mcr*-1 prevalence was 68.86 % (303/440) in pigs, 87.58 % (388/443) in chicken, and 71.43 % (30/42) in cattle, compared with *mcr*-2 prevalence of 46.82 % (206/440) in pigs, 14.90 % (66/443) in chicken, and 19.05 % (8/42) in cattle. The *mcr* variant gene prevalence reported by [17] was considerably higher than ours and those previously reported in China which was based on the presence of the *mcr* in bacterial isolates. They directly detected the clinical samples instead of isolated *E.coi* strains and sequenced three variants of *mcr*-1, *mcr*-2, and *mcr*-3 [17]. The *mcr-1* and *mcr-2* occurred widely in pigs and poultry of Chinese farms [17–18]. Except harbouring the *mcr* genes, a *mcr*-independent mechanism behind the remaining colistin resistant *E.coli* colonies, for example, lipopolysaccharide modification [19], other (transferable) colistin-resistant mechanisms, and undefined mechanisms exist. The implication of the *mcr* gene wide spread will be enormous if plasmid-mediated colistin resistance was readily passed between *E. coli* strains, and also be passed to *Klebsiella pneumoniae* and *Pseudomonas aeruginosa* strains like descriptions in *the Lancet Infectious Disease* published by Liu Yi-Yun and colleagues [2]. Since 1 April 2017, the Chinese government has implemented the withdrawal of colistin as a food additive for growth promotion in food animal [20], this policy is in line with international policy of One Health.

## 5. CONCLUSION

The management of colistin resistance at the human-animal-environment interface requires the urgent use of the One Health approach for effective control and prevention. Our study will provide new data about colistin resistance prevalence worldwide. The colistin resistance widely occurred at all ages of domestic animals and the *mcr*-dependent mechanism dominated in *E.coli*. We also found that the older and adult animals were a reservoir of resistant strains, suggesting a potential food safety issue and greater public health problems.

## AUTHOR CONTRIBUTIONS

Zhang XH and He KW conceived and designed the experiments. Zhang BC analyzed the data. Yu ZY, GuoYY, WangJ, Zhao PD, and Liu JJ performed the experiments; Zhang XH and Zhang BC wrote the paper.

## Ethical approval

Ethical approval was granted by the Ethics Committee of the Institute of Veterinary Medicine, Jiangsu Academy of Agricultural Sciences (Nanjing, Jiangsu, China) [20150212].

## REFERENCES

[1] Investigation of *E. coli* Harboring the *mcr-* 1Resistance Gene — Connecticut, 2016. Morbidity and Mortality Weekly Report (MMWR) 2016; 65(36), 979–80.

[2] Liu YY, Wang Y, Walsh TR, Yi LX, Zhang R, Spencer J, Doi Y, Tian G, Dong B, Huang X, Yu LF, Gu D, Ren H, Chen X, Lv L, He D, Zhou H, Liang Z, Liu JH, Shen J. 2016. Emergence of plasmid-mediated colistin resistance mechanism MCR-1 in animals and human beings in China: a microbiological and molecular biological study. The Lancet Infect Dis 16(2): 161–8.

[3] Nicolet S, Goldenberger D, Schwede T, Page M, Creus M. 2016. Plasmid-mediated colistin resistance in a patient infected with *Klebsiella* pneumoniae. The Lancet Infect Dis 16(9): 998–9.

[4] Van Boeckel TP, Brower C, Gilbert M,, Grenfell BT, Levin SA, Robinson TP, Teillanta A, Laxminarayan R. 2015. Global trends in antimicrobial use in food animals. Proc Natl Acad Sci U S A 112(18): 5649–54.

[5] Guyonnet J, Manco B, Baduel L, Kaltsatos V, Aliabadi MH, and Lees P. 2010. Determination of adosage regimen of colistin by pharmacokinetic/pharmacodynamics integration and modeling for treatment of G.I.T. disease in pigs. Res Vet Sci 88, 307–14.

[6] Rhouma M, Beaudry F, and Letellier A. 2016. Resistance to colistin: what is the fate for this antibiotic in pig production? Int J Antimicrob Agents 48, 119–26.

[7] Miyagu chi H, Narita H, Sakamoto K. 1996. An antibiotic-binding motif of an RNA fragment derived from the A site related region of *E. coli* 16S rRNA. Nucleic Acids Res 24: 3700–6.

[8] Xavier BB, Lammens C, Ruhal R, Kumar-Singh S, Butaye P, Goossens H, Malhotra-Kumar S. 2016. Identification of a novel plasmid-mediated colistin-resistance gene, *mcr-2*, in *Escherichia coli*, Belgium, June 2016. Euro Surveill 21 (27).

[9] Hoelzer K, Wong N, Thomas J, Talkington K, Jungman E, Coukell A. 2017. Antimicrobial drug use in food-producing animals and associated human health risks: what, and how strong, is the evidence? BMC Vet Res 13(1):211.

[10] China Veterinary Drug Association. Annual Report on Development of Veterinary Medicine Industry in China (2012). Beijing: China Agriculture Press 2014.

[11] Jans C, Sarno E, Collineau L, Meile L, Stärk KDC, Stephan R. 2018. Consumer exposure to antimicrobial resistant bacteria from food at Swiss retail level. Front Microbiol 9:362.

[12] Liebana E, Carattoli A, Coque TM, Hasman H, Magiorakos AP, Mevius D, Peixe L, Poirel L, Schuepbach-Regula G, Torneke K, Torren-Edo J, Torres C, Threlfall J. 2013. Public health risks of enterobacterial isolates producing extendedspectrum beta-lactamases or AmpC beta-lactamases in food and food producing animals: an EU perspective of epidemiology, analytical methods, risk factors, and control options. Clin Infect Dis 56, 1030–7.

[13] Huang X, Yu L, Chen X, Zhi C, Yao X, Liu Y, Wu S, Guo Z, Yi L, Zeng Z, Liu J. 2017. High prevalence of colistin resistance and mcr-1 gene in *Escherichia coli* isolated from food animals in china. Front Microbiol 8:562.

[14] Rhouma M, Beaudry F, Thériault W, Letellier A. 2016. Colistin in Pig Production: Chemistry, Mechanism of Antibacterial Action, Microbial Resistance Emergence, and One Health Perspectives. Front Microbiol 11:7.

[15] Rhouma M, Beaudry F, Thériault W, Bergeron N, Laurent-Lewandowski S, Fairbrother JM, Letellier A. 2015. Gastric stability and oral bioavailability of colistin sulfate in pigs challenged or not with *Escherichia coli* O149: F4(K88). Res Vet Sci 102, 173–81.

[16] Fukuda A, Sato T, Shinagawa M, Takahashi S, Asai T, Yokota S-i, Usui M, Tamura Y. 2018. High prevalence of *mcr-1, mcr-3*, and *mcr-5* in *Escherichia coli* derived from diseased pigs in Japan. Int J Antimicrob Agents 51(1): 163–4.

[17] Zhang J, Chen L, Wang J, Yassin AK, Butaye P, Kelly P, Gong J, Guo W, Li J, Li M, Yang F, Feng Z, Jiang P, Song C, Wang Y, You J, Yang Y, Price S, Qi K, Kang Y, Wang C. 2018. Molecular detection of colistin resistance genes (mcr-1, *mcr-2* and *mcr-3)* in nasal/oropharyngeal and anal/cloacal swabs from pigs and poultry. Sci Rep27, 8(1):3705.

[18] Yassin AK, Zhang J, Wang J, Chen L, Kelly P, Butaye P, Lu G, Gong J, Li M, Wei L, Wang Y, Qi K, Han X, Price S, Hathcock T, Wang C. 2017. Identification and characterization of *mcr* mediated colistin resistance in extraintestinal *Escherichia coli* from poultry and livestock in China. FEMS Microbiol Lett 29, 364(24).

[19] Guerin F, Isnard C, Sinel C, Morand P, Dhalluin A, Cattoir V, Giard JC. 2016. Cluster-dependent colistin hetero-resistance in Enterobacter cloacae complex. J Antimicrob Chemother 71(11): 3058–61.

[20] Wang X, Liu Y, Qi X, Wang R, Jin L, Zhao M, Zhang Y, Wang Q, Chen H, Wang H. 2017. Molecular epidemiology of colistin-resistant Enterobacteriaceae in inpatients and avian from China: high prevalence of mcr-negative *Klebsiella pneumoniae*. Int J Antimicrob Ag 50(4):536–41.

